# Functional properties of habenular neurons are determined by developmental stage and sequential neurogenesis

**DOI:** 10.1101/722462

**Authors:** Stephanie Fore, Mehmet Ilyas Cosacak, Carmen Diaz Verdugo, Caghan Kizil, Emre Yaksi

## Abstract

Neural development is not just a linear expansion of the brain. Instead, the structure and function of developing brain circuits undergo drastic alterations that have a direct impact on the animals’ expanding behavioural repertoire. Here we investigated the developmental changes in the habenula, a brain region that mediates behavioural flexibility during learning, social interactions and aversive experiences. We showed that developing habenular circuits exhibit multiple alterations, which increase the structural and functional diversity of cell types, inputs and functional modules within habenula. As the neural architecture of habenula develops, it sequentially transforms into a multi-sensory brain region that can process visual and olfactory information. Moreover, we also observed that already at early developmental stages, the habenula exhibits spatio-temporally structured spontaneous neural activity that shows prominent alterations and refinement with age. Interestingly, these alterations in spontaneous activity are accompanied by sequential neurogenesis and integration of distinct neural clusters across development. Finally, by combining an in vivo neuronal birthdating method with functional imaging, we revealed that clusters of habenular neurons with distinct functional properties are born sequentially at distinct developmental time windows. Our results highlight a strong link between the function of habenular neurons and their precise birthdate during development, which supports the idea that sequential neurogenesis leads to an expansion of neural clusters that correspond to distinct functional modules in the brain.

## INTRODUCTION

During development and maturation, the functional capacity of brain circuits increases in order to support the animals’ expanding behavioural repertoire. For example, while a young zebrafish larvae mostly needs to avoid threats (Burgess and Granato, 2007; Heap et al., 2018; Temizer et al., 2015) and find food (Lindsay and Vogt, 2004), more cognitively demanding behaviours such as associative learning (Valente et al., 2012) or social interactions (Dreosti et al., 2015; Hinz and de Polavieja, 2017) emerge later in development during the juvenile stage around 3-4 weeks. Such expansion of the behavioural repertoire is a feature that is conserved across vertebrates (Anderson, 1984; Brenowitz and Beecher, 2005), and often accompanied with the development, establishment and maturation of distinct circuit components generating functional modules within the brain (Bjerknes et al., 2018; Donato et al., 2017; Tottenham and Gabard-Durnam, 2017). Hence, brain development and maturation is not just a linear expansion of the existing building blocks, but also a sequential increase in the diversity of functional modules and cell types. In the cortex for example, individual layers are born at different time points, with deep layers being born earlier than superficial layers (Angevine and Sidman, 1961; Lein et al., 2017; Luskin and Shatz, 1985), creating a diversity in the cytoarchitecture of cortical layers, cell types and regions with distinct functions (Donato et al., 2017; Jabaudon, 2017; Sur and Leamey, 2001). The maturation and refinement of the developing brain relies on neural activity, which can be evoked by sensory inputs (Penn and Shatz, 1999; Wiesel and Hubel, 1965), or spontaneously generated (Galli and Maffei, 1988; Moreno-Juan et al., 2017; O’Donovan, 1989; Penn and Shatz, 1999; Yuste, 1997). In most sensory systems, sensory evoked activity was shown to be critical for the maturation of neural circuits and the refinement of topographical maps (Katz and Shatz, 1996; Webster and Webster, 1977; White et al., 2001; Wiesel and Hubel, 1965; Zou et al., 2004). The appearance of spontaneous activity of the developing brain starts early, and coincides with periods of intense synaptogenesis and neuronal growth (Ben-Ari, 2001; Galli and Maffei, 1988; Maffei and Galli-Resta, 1990). For example, spontaneous activity bursts were observed in the visual, auditory and somatosensory systems (Elstrott et al., 2008; Feller, 1999), and are shown to be important for the remodelling of these structures (Katz and Shatz, 1996; Moreno-Juan et al., 2017; Xu et al., 2011). In higher brain regions, such as the hippocampus, large and slow bursts are observed (Leinekugel et al., 1998; Rudy et al., 1987) before the appearance of faster rhythms and the patterned activity of the adult hippocampus associated with learning and memory (Altman et al., 1973; Rudy et al., 1987). Moreover, spontaneous activity is also important for the maturation of connections across distant brain regions, as observed between the olfactory bulb and the entorhinal cortex (Gretenkord et al., 2019). Finally, the synchronous bursting during spontaneous activity is shown to be mediated via excitatory connections (Meister et al., 1991; O’Donovan et al., 1998), gap junctions (Yuste et al., 1992), glia (Zhang et al., 2019) or other support cells (Babola et al., 2018), and plays an important role in establishing neural connectivity (Kerschensteiner, 2014; Yu et al., 2012). Taken together, both sensory-driven and spontaneous neural activity are essential in the development and maturation of neural circuits. It is however less clear how sensory and spontaneous activity dependent processes interact, and shape each other as the animals develops.

While the majority of studies have focused on studying activity-dependent processes in the development of sensory systems, less is known about the maturation of higher brain areas integrating information from multiple brain regions, such as the habenula. The habenula is a particularly interesting brain region, as it was shown to integrate both sensory (Cheng et al., 2017; Miyasaka et al., 2009; Zhang et al., 2017) and cortico-limbic inputs (Herkenham and Nauta, 1977; Hong et al., 2011; Lazaridis et al., 2019; Matsumoto and Hikosaka, 2007; Meye et al., 2016; Okamoto et al., 2012; Stamatakis et al., 2013; Turner et al., 2016; Warden et al., 2012) while directly regulating the function of monoaminergic brain nuclei controlling behaviour (Duboue et al., 2017; Flanigan et al., 2017; Hikosaka, 2010; Krishnan et al., 2014; Lin and Jesuthasan, 2017; Matsumoto and Hikosaka, 2007; Zhang et al., 2017). Dysfunction of the habenula is also shown to be associated with several neurological conditions and mood disorders including depression(Huang et al., 2019; Yang et al., 2018). The habenula is composed of several subdomains or modules based on its neurochemical profiles (Aizawa et al., 2012; deCarvalho et al., 2014; Pandey et al., 2018; Ren et al., 2011; Seigneur et al., 2018; Viswanath et al., 2013), anatomical projections (Agetsuma et al., 2010; Amo et al., 2010; Amo et al., 2014; Lee et al., 2010), and the activity of habenular neurons (Amo et al., 2014; Dreosti et al., 2014; Jetti et al., 2014). Two major subnuclei, the lateral and medial habenula in mammals, respectively dorsal habenula (dHb) and ventral habenula (vHb) in zebrafish, have clear differences in their functional and molecular profiles (Amo et al., 2014; Lee et al., 2010). While the dHb is involved in sensory processing (Lin and Jesuthasan, 2017; Zhang et al., 2017), circadian rhythms (Lin and Jesuthasan, 2017), aggression (Chou et al., 2016) and experience dependent fear response (Agetsuma et al., 2010; Duboue et al., 2017; Lee et al., 2010), the vHb was shown to play a role in avoidance learning (Amo et al., 2014) and active coping behaviour (Andalman et al., 2019). In zebrafish, dHb and vHb originate from separate neural progenitor pools, and were suggested to maturate at distinct developmental time points (Beretta et al., 2013). Moreover, the dHb was shown to undergo asymmetric neurogenesis during the early stages of development, with left dHb neurons being born earlier than the ones in the right hemisphere (Aizawa et al., 2007). The prominent asymmetry in the molecular (deCarvalho et al., 2014) and anatomical properties of habenula (Ahumada-Galleguillos et al., 2017; Bianco et al., 2008), is also reflected in its asymmetric encoding of visual and olfactory information in the left and the right dHb hemispheres respectively (Dreosti et al., 2014; Jetti et al., 2014; Miyasaka et al., 2009). A recent transcriptome analysis further revealed multiple molecularly distinct and spatially organized functional modules within the zebrafish habenula (Pandey et al., 2018), which resemble the functional clusters of habenular neurons that are spontaneously active (Jetti et al., 2014). All these evidences, further suggests a fine spatial organization of distinct functional modules within habenula (Jetti et al., 2014). In addition, spontaneous habenular activity was shown to govern sensory responses and therefore proposed to represent internal states of the network, which could mediate the selection of appropriate behaviours (Jetti et al., 2014). In fact, a recent study revealed a sequential recruitment of neurons generating an increase in vHb activity, during the switch from active to passive coping behaviours in juvenile zebrafish (Andalman et al., 2019). Interestingly, such complex behaviours emerge at later stages of zebrafish development (Dreosti et al., 2015; Hinz and de Polavieja, 2017; Valente et al., 2012; Yashina et al., 2019), suggesting pronounced changes in the underlying circuitry across the brain. Despite extensive characterization of habenular circuitry during very early developmental stages (Aizawa et al., 2007; Bianco et al., 2008; deCarvalho et al., 2014), it is still unclear how the maturation of habenular networks and the establishment of its distinct functional modules corresponds to the observed expansion of animals behavioural repertoire during development.

In the present study, we investigated how the function and the architecture of habenular networks changes across development, and how these alterations relate to the formation of distinct functional modules within habenula. We observed that as the habenula expands, the number of neurons increases, inhibitory connections are formed and sensory inputs are integrated with a temporal order. We showed that visual responses are more prominent early on, but olfactory responses quickly catch up, while both sensory modalities are distinctly encoded in habenula at all developmental stages. Spontaneous habenular activity is present even at early developmental stages and it is strongly predictive of the sensory responses in habenula. Interestingly, we observed a prominent restructuring of both spatial and temporal features of spontaneous habenular activity during development. These changes in the habenula were accompanied by a sequential and spatially organized addition of newly born neurons across development. Finally, we developed an in vivo birthdating method and directly showed that habenular neurons born at a distinct developmental stage, exhibit highly correlated spontaneous activity with similar functional features. Taken together, these results revealed that distinct functional clusters of habenular neurons are born at different developmental stages. Our observations support the idea that neuronal birthdate and function are strongly related. We propose that this functional refinement of the habenular circuits underlies the transition of a developing zebrafish larvae into a mature animal with increased cognitive capacity.

## RESULTS

### New neurons and new inhibitory connections are added to habenula during development

To study the functional development of zebrafish habenula, we focused on three developmental stages: 3dpf (early larval stage), 6dpf (late larval stage) and 21dpf (juvenile stage). During these periods, zebrafish transition from self-feeding larvae (via the yolk sac) to an external feeding juvenile animal with more complex behaviours including those involving habenular circuits (Agetsuma et al., 2010; Amo et al., 2014; Andalman et al., 2019; Dreosti et al., 2015; Hinz and de Polavieja, 2017; Valente et al., 2012). First, we visualized the gross morphology of Hb in Tg(elavl3:GCaMP6s) (Vladimirov et al., 2014) zebrafish, labelling most habenular neurons. We observed that the number of habenular neurons increases drastically from 3 to 21 days of zebrafish development (Figure 1A-B). Next we investigated the expression of glutamate and GABA by using Tg(vglut2a:dsRed) (Miyasaka et al., 2009) and Tg(gad1b:GFP) (Satou et al., 2013) zebrafish. We found that at 3 days, habenula mainly consists of glutamatergic neurons, and as the animal get older few GABAergic neurons are added (Figure 1C). Interestingly, from 6dpf on we also observed prominent bilateral GABAergic projections that innervate specific domains at the lateral sides of habenula (Figure 1C). These results suggest that habenula undergoes a non-linear expansion with the addition of new cell types and inputs arriving at different time points throughout development. This could potentially lead to the formation of new functional modules within habenula.

**Figure 1:**
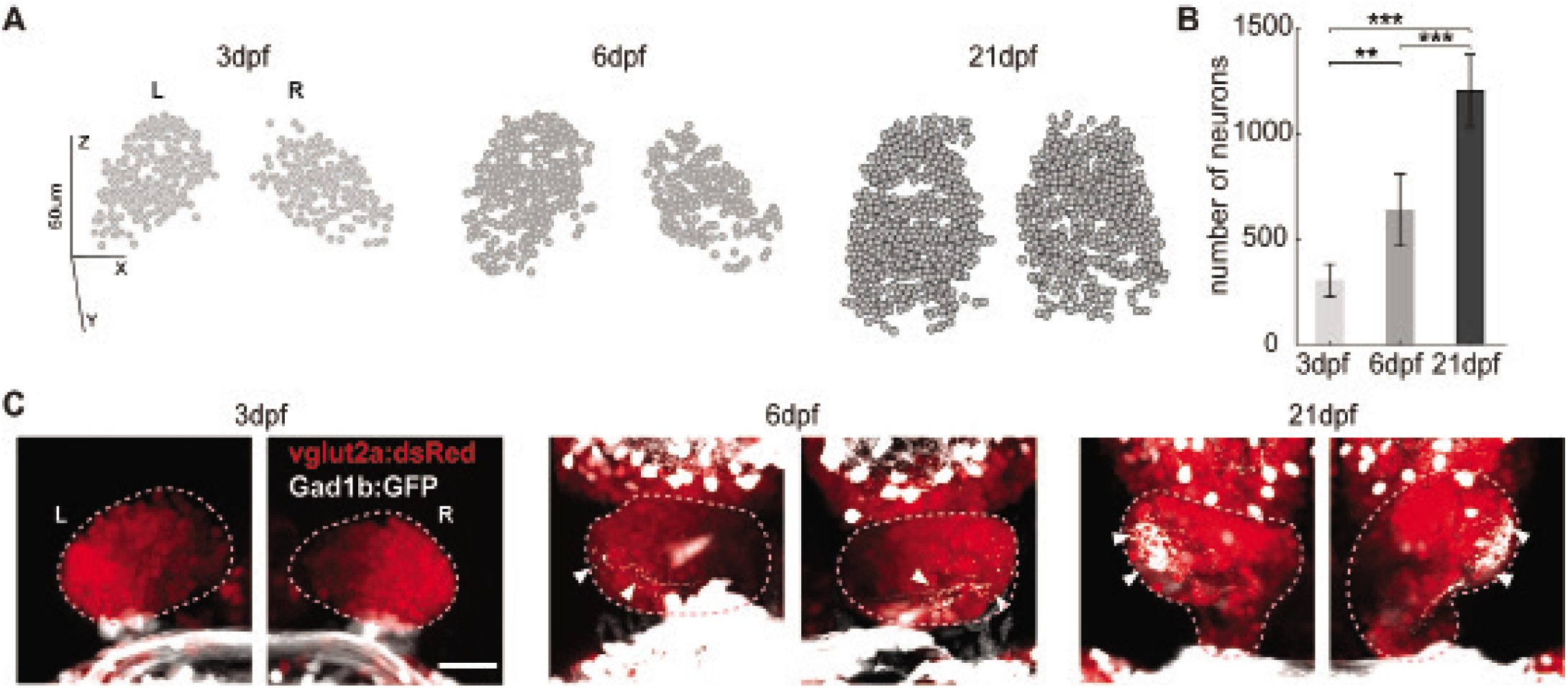
During habenular development the number of glutamatergic neurons increases and new GABAergic connections are added. (A) Representative examples of three-dimensional reconstructions of habenular neurons detected in Tg(elavl3:GCaMP6s) zebrafish line, using volumetric two-photon calcium imaging at three different developmental ages (3, 6 and 21dpf). Coronal view. (B) The average number of neurons detected in Tg(elavl3:GCaMP6s) at 3dpf (n=9 fish), 6dpf(n=6 fish) and 21dpf (n=9 fish), **p<0.01, ***p<0.001, t-test. Data presented as mean ±SEM. (C) Confocal microscopy projections of habenula in Tg(vglut2a:dsRed) and Tg(gad1b:GFP) double transgenic zebrafish at 3,6 and 21dpf, labelling glutamatergic (red) and GABAergic (white) neurons and projections, dorsal view. White arrows indicate GABAergic projections. White scale bar represents 100μm. L,left; R,right hemisphere.

### Sensory inputs and computations in habenula follow a developmental order

Previous studies showed that habenula receives olfactory (Miyasaka et al., 2009) and visual (Cheng et al., 2017; Zhang et al., 2017) inputs and responds to both of these sensory modalities (Dreosti et al., 2014; Jetti et al., 2014; Krishnan et al., 2014). To investigate how these sensory inputs maturate during habenula development, we visualized the mitral cell and thalamic axon terminals in habenula using Tg(lhx2a:gap-YFP) (Miyasaka et al., 2009) and Et(−0.6hsp70l:Gal4-VP16) s1020t; Tg(UAS:nfsB-mCherry) (Agetsuma et al., 2010; Mohamed et al., 2017; Scott and Baier, 2009) zebrafish lines, respectively. At all ages, thalamic projections were found to innervate habenula (Figure 2A, blue) in distinct locations that are different from GABAergic innervations (Figure 2A, white). Yet, the density of olfactory bulb projections increased with age (Figure 2A, red). These findings suggest that habenula neurons are predominantly light responsive at younger ages, while odour responses develop later. In order to test this hypothesis, we measured odour (food odour) and light (red light flash) responses in the habenula with two-photon calcium imaging using Tg(elavl3:GCaMP6s) (Vladimirov et al., 2014) zebrafish. At 3 days we found that a significantly higher portion of habenular neurons is responsive to the visual stimulus compared to odour stimulus (Figure 2B-C). As the animals develop, the ratio of odour and light responding neurons became similar, suggesting a shift from a predominantly visual to a multi-sensory processing habenula at older ages. It is also worth noting that both visual and olfactory responses display faster decays in the older animals (Figure 2B, Supplementary figure 1A), which might be due to the increased inhibition at the older stages (Figure 1C). Next, we investigated how visual and olfactory information are encoded in the activity of habenular neurons. We found that already at younger ages distinct populations of habenular neurons preferentially responded to either olfactory or visual stimuli (Figure 2D). More specifically, the correlations between the olfactory and visual responses remained low at all developmental stages (Figure 2D, inserts), suggesting that the habenular neurons are selective for different sensory modalities from early on and persists across development. These results highlight the importance of olfactory and visual information for the functioning of habenular networks and related behaviours already at the early stages of habenula development.

**Figure 2:**
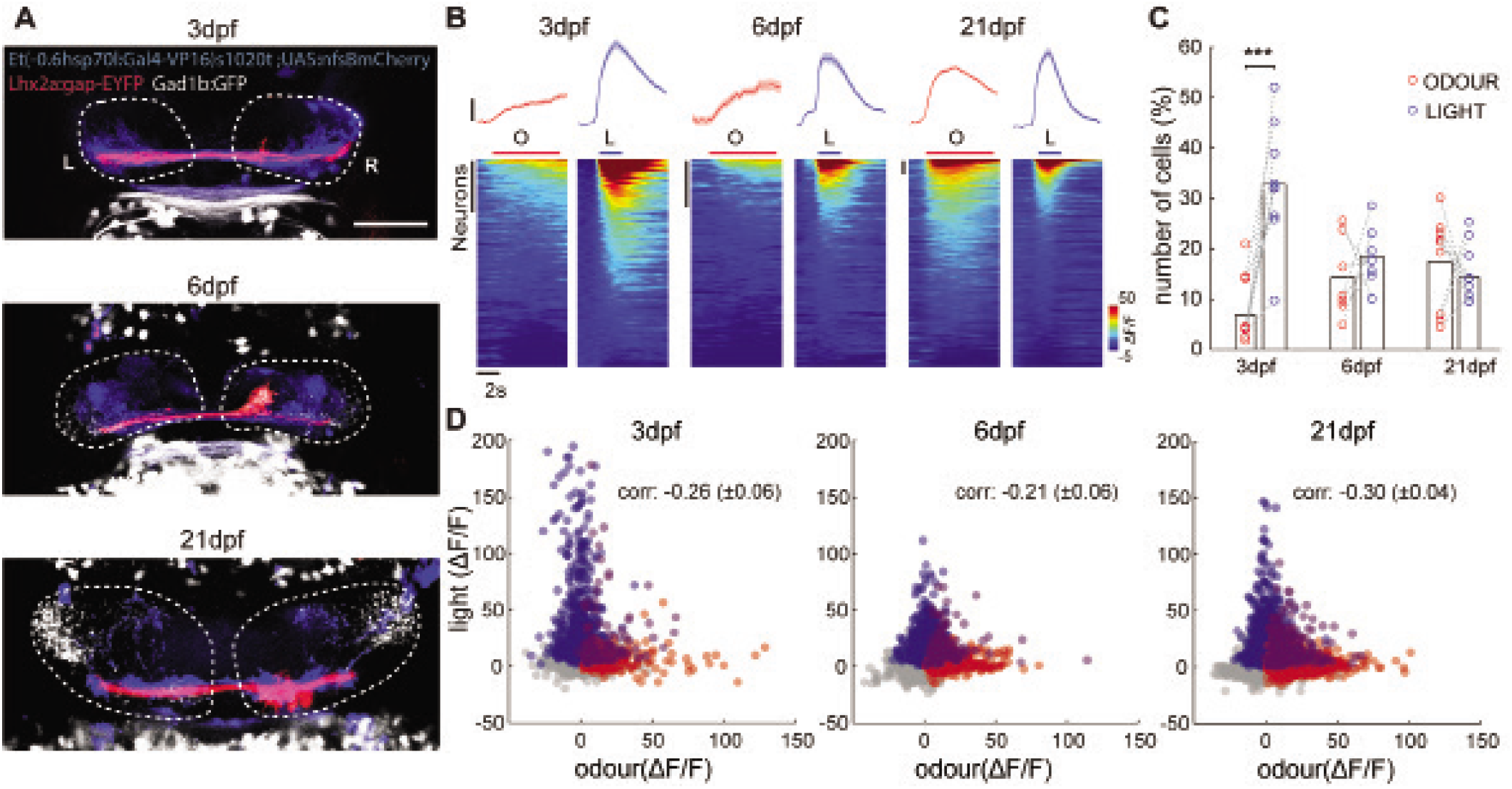
Visual responses dominate larval habenula, while olfactory responses develop later. (A) Confocal microscopy projections of habenula in Tg(Lhx2:gap-YFP); (Et(0.6hsp70l:Gal4-VP16) s1020t; Tg(UAS:nfsB-mCherry) and Tg(gad1b:GFP) triple transgenic zebrafish labelling olfactory bulb projections (blue), thalamic projections (red) and GABAergic projections (white), dorsal view. Dashed white lines delineate habenula. White bar represents 100μm. L,left; R,right hemisphere. (B) Top panel shows average responses of odour-responding (red) and light-responding habenular neurons (blue) over 4-5trials in at 3dpf (n=9), 6dpf (n=7) and 21dpf (n=9) zebrafish. Black bars represent 10% ΔF/F. Shadows represent SEM. Bottom panel shows color coded sensory responses (ΔF/F) of all individual Hb neurons exposed to 5s odour stimulus (left) and 2s light stimulus (right) within a representative fish imaged at 3,6 and 21dpf. Black bars represent 100 neurons. (C) Percentage of odour (red dots) vs light (blue dots) responding neurons at 3,6 and 21dpf. ***p<0.001 paired t-test. (D) Odour (red) vs light (blue) response amplitude of all habenular neurons responses at 3,6 and 21dpf. Magenta color depicts the neurons responding to both odour and light. “corr” represents the mean correlation value of neural responses to odour vs light responses averaged across all fish ±SEM. Note that at all ages odour and light responses are dissimilar, depicted by the negative correlations.

### Functional lateralization of sensory responses is a feature of dorsal habenula, but not prominent in ventral habenula that develops later

Earlier studies revealed a strong molecular (Amo et al., 2010; deCarvalho et al., 2014; Lee et al., 2010; Viswanath et al., 2013) and functional (Dreosti et al., 2014; Krishnan et al., 2014) lateralization of habenular networks especially in the dHb of larval zebrafish. Interestingly, there is no evidence for such lateralization in the rodent habenula, which is mostly studied during adult stages. This made us question whether the ontogeny of functional lateralization in habenula might resemble the phylogeny of the vertebrate habenula, where asymmetric distributions of neural populations appear first and symmetric distributions develop later. To test this hypothesis we investigated the lateralization of visual and olfactory responses in habenular neurons across development (Figure 3). We observed that the dorsal sections of habenula, which likely covers mostly the dHb, showed lateralized sensory responses at all ages (Figure3A). We quantified this functional lateralization by measuring the ratio of sensory responding neurons across hemispheres (Dreosti et al., 2014) and by using a lateralization index, where 0 represent full symmetry and 1 represent a complete segregation of sensory responses across habenular hemispheres. We observed that dHb exhibit high functional lateralization (Figure 3B-D). This contrasts to the symmetric sensory responses in the ventral sections of habenula, with low lateralization index across development (Figure 3B-D). We also observed a decrease in sensory responding neurons in the ventral sections of habenula especially at 21 dpf (supplementary Figure 1B). Despite the reduced functional lateralization, we observed a prominent spatial organization of sensory responses within habenular hemispheres. We quantified this spatial organization by using the measure of focality index, ranging from 0 (for random spatial distribution of sensory responses) to 1 (for maximally focal sensory responses) (Yaksi et al., 2007). At all ages we observed that the focality index for sensory responding neurons was non-random and significantly higher than non-responding neurons (Figure 3E). These results indicate that while sensory responses are spatially organized from the early larval stages, the lateralization of sensory computations are mainly a feature of dHb (Dreosti et al., 2014) that develops earlier. Instead, vHb, which develops later during development, exhibits fewer sensory responses and decreased lateralization.

**Figure 3:**
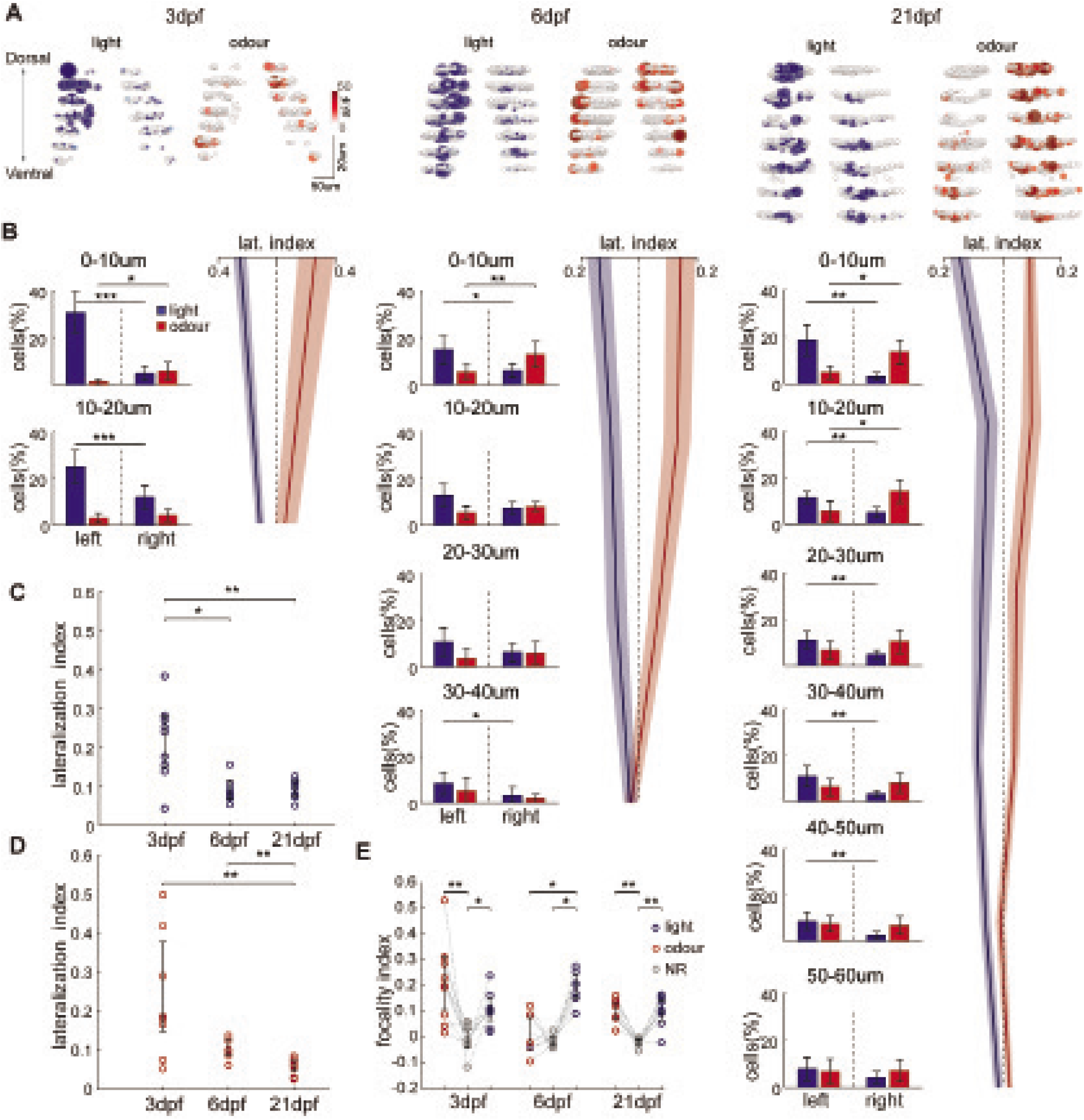
Sensory lateralization is prominent in dorsal habenula at all ages, but decreases with age especially in ventral habenula. (A) Three-dimensional reconstructions of habenular responses to light (blue) and odour (red) stimulation recorded by volumetric two-photon microscopy in Tg(elavl3:GCaMP6s) zebrafish, at 3,6 and 21dpf. White dots indicates non-responding neurons. Coronal view. (B) The ratio of odour (red) vs light (blue) responding neurons in left vs right habenular hemispheres, as well as associated sensory lateralization index at 3 (left), 6 (middle) and 21 (right) dpf zebrafish. Each row represent the average values for neurons within 10μm plane (from the top to bottom). Shades represent the ±SEM. 3dpf (n=9 fish), 6dpf (n=7 fish) and 21dpf (n=9 fish). *p<0.05, **p<0.01, ***p<0.001, t-test. (C) Average lateralization index of light responses for the entire habenula of all fish at 3,6,21dpf. *p<0.05, **p<0.01, Wilcoxon ranksum test. (D) Average lateralization index of odour responses for the entire habenula of all fish at 3, 6, 21dpf. **p<0.01, Wilcoxon ranksum test.

### Spatially organized functional clusters of habenular neurons exhibit spontaneous activity and are recruited during sensory stimulation at all ages

In juvenile zebrafish, spatially organized functional clusters of habenular neurons exhibit synchronous bursts of spontaneous activity, which is present even in the absence of sensory inputs (Jetti et al., 2014). It is however less clear if such spontaneous activity is also present in younger zebrafish larvae, and to what extent this endogenously generated activity relates to the maturation of habenular neurons. To study such processes, we recorded the ongoing activity in habenula in the absence of external stimuli at different developmental stages described above. We observed that spontaneous activity is present from 3dpf and functional clusters of habenular neurons exhibit correlated activity (Figure 4A). Next, we investigate the stability of these functional clusters during different time windows, using k-means clustering. We showed that at all developmental stages, a large portion of habenular neurons that belong to a functional cluster remain in their respective cluster with a significantly higher probability compared to chance levels (Figure 4B). To investigate the synchrony between the habenular neurons further, we next computed the ratio of significantly positive and negative correlated pairs of habenular neurons. We observed that as the animals develops, more pairs of neurons displayed significant and robust positive correlations (Figure 4C). However, we did not observe such a change for significant negative correlations between habenular neurons (Figure 4D). Next, we investigated the spatial organization of synchronous habenular activity by plotting the average pairwise correlation of neurons versus the distance between them. We observed that while at all developmental stages nearby habenular neurons have more correlated spontaneous activity, the nearby habenular neurons exhibit significantly stronger correlations during older developmental stages (Figure 4E). Such increase in the ratio (Figure 4C) and strength (Figure 4E) of positive correlations, is likely due to maturation of synaptic connections between habenular neurons as well as synaptic inputs to habenula.

**Figure 4:**
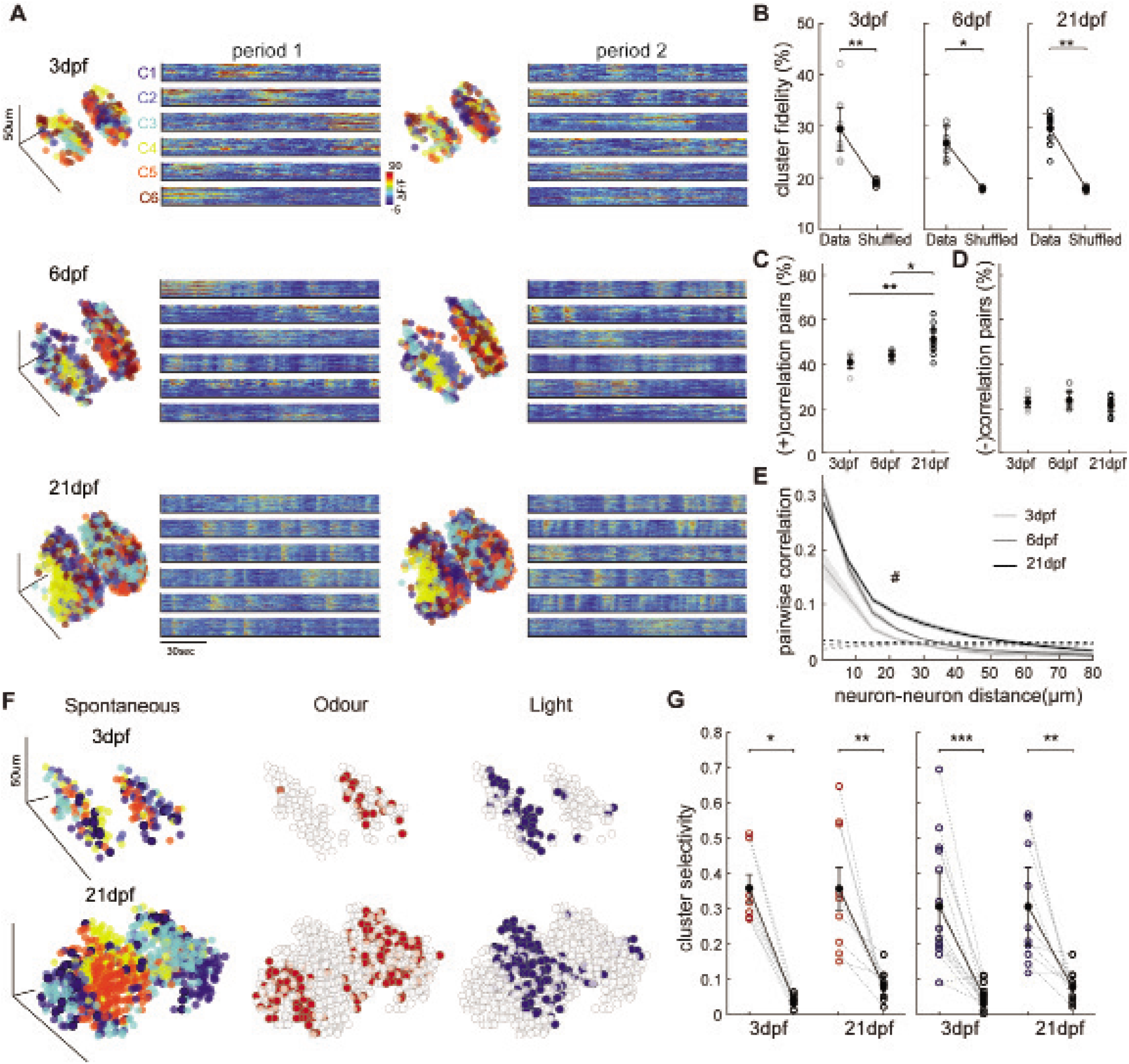
Spontaneous habenular activity is structured into spatially organized functional clusters of neurons that are preserved during sensory responses, at all ages. (A) Functional clusters of habenular neurons with synchronous spontaneous activity in two different time periods (period 1 and period 2). Three-dimensional reconstruction depicts the spatial locations of all neurons. Each colour represents a neuronal cluster with similar spontaneous activity, defined by using k-means clustering. Colour coded neural traces on the right, show the spontaneous activity of each neuron that belongs to a given cluster “C”. warm colours represents increased neural activity. Each row represents an example zebrafish at 3, 6, 21dpf. (B) Cluster fidelity, depicting the ratio of neural pairs that remain within the same functional cluster during 2 different time periods for 3dpf (n=9), 6dpf (n=6) and 21dpf (n=9), compared to shuffled chance levels. *p<0.05, **p<0.01, Wilcoxon signed-rank test. (C) The ratio of neural pairs with significant (p<0.05) positive correlations during two periods of spontaneous activity. *p<0.05, **p<0.01, Wilcoxon ranksum test. (D) The ratio of neural pairs with significant (p<0.05) negative correlations during two periods of spontaneous activity. Wilcoxon ranksum test. (E) Relation between pairwise correlation of spontaneous neural activity and the distance between each neuron pair in habenula at 3, 6, 21dpf. Dotted lines represent the results when neuron locations are shuffled. ANOVA-n displayed significance over distances (p<0.001) and over age groups (p<0.001) indicated with #. (F) Colour coded functional clusters of habenular neurons with synchronous spontaneous activity for an example zebrafish at 3 (top) and 21(bottom) dpf. On the middle and left panels, odour (red) and light (blue) responses of the same neurons are visualized. Scale bars represent 50μm. (G) Cluster selectivity, depicting how odour (red) or light (blue) responding neurons are distributed into functional clusters based on their spontaneous activity. High selectivity means that both odour and light responding neurons belong to fewer clusters than non-responding neurons (grey dots), at 3 and 21dpf. *p<0.05, **p<0.01, ***p<0.001, Wilcoxon signed-rank test. All data presented as mean ±SEM.

What is the function of spontaneous activity in habenula at early developmental stages? In the visual system for example, spontaneous activity waves are important for establishing the prior patterning of eye specific connections before the onset of sensory input (Penn and Shatz, 1999). To investigate this possibility, we investigated whether spontaneous habenular activity is related to sensory responses in habenula, and if those neurons with correlated spontaneous activity also respond to sensory stimulation similarly. We showed that already at the early developmental stages, odour and light responsive habenular neurons exhibit distinct patterns of spontaneous activity and largely overlap with the distinct functional clusters within habenula. (Figure 4F-G). We observed that the relationship between the spontaneous and sensory evoked activity of habenular neurons remains stable across development, highlighting the importance of spontaneous activity in predicting the sensory response characteristics of habenular neurons.

### Temporal features of spontaneous habenular activity change during development as new neurons are sequentially arranged into distinct spatial domains

Our results revealed that the spontaneous activity is a feature of habenula already at early developmental stages. In several brain regions spontaneous activity during early development is generally characterized as large bursts with long quiescent intervals (Elstrott et al., 2008; Feller, 1999; Leinekugel et al., 1998), whereas adult brains usually exhibit rhythmic activity at faster frequencies (Buzsaki and Draguhn, 2004; Leinekugel et al., 1998). To test whether the temporal features of habenular activity are changing over development, we detected the spontaneous activity bursts (Romano et al., 2017) (Figure 5A) and quantified the frequency and duration of these events. We observed that as the animals develop, the frequency of spontaneous activity bursts significantly increases (Figure 5B) and their duration decreases (Figure 5C). Moreover, while the average activity of each habenular neuron decreases (Figure 5D), a larger proportion of habenular neurons exhibits spontaneous activity(Figure 5E) at later developmental stages. These results are in line with the developmental changes observed in other vertebrate brain regions and highlight the maturation of habenular spontaneous activity across development.

**Figure 5:**
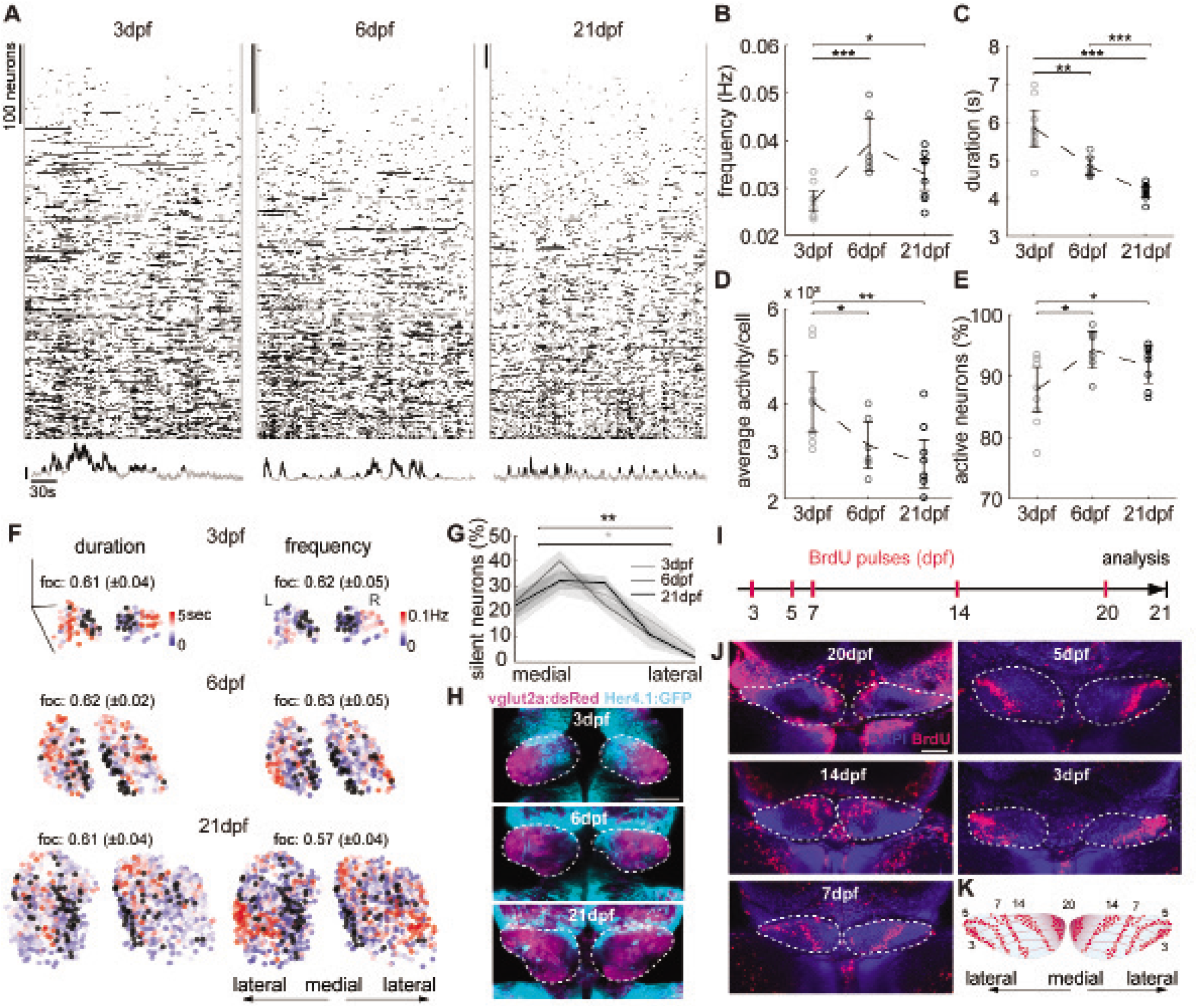
Temporal features of habenular spontaneous activity change during development as new neurons are sequentially arranged into distinct spatial domains. (A) Individual spontaneous activity bursts (black markings) detected in all habenular neurons at 3,6 and 21dpf. Bottom panel shows individual example spontaneous activity traces from three different neurons, with detected events marked with black colour. Black bar represent 10% ΔF/F. (B) Average frequency of spontaneous activity bursts in habenular neurons in each zebrafish at 3dpf (n=9 fish), 6dpf (n=6 fish) and 21dpf (n=9 fish). (C) Average duration of spontaneous activity bursts in habenular neurons in each zebrafish. (D) Average spontaneous activity in habenular neurons in each zebrafish, represented as total area under the curve of all detected events. (E) The ratio of spontaneously active neurons with at least one spontaneous activity burst.*p<0.05, **p<0.01, ***p<0.001,Wilcoxon ranksum test. (F) Three-dimensional reconstructions of habenular neurons based on the duration (left) and frequency of their spontaneous activity. Frequency and duration are colour coded. Black dots are inactive neurons. Coronal view. Scale bars represent 50μm. ‘foc’ represents the focality index as mean ±SEM for the temporal features of spontaneous activity bursts, duration and frequency, across all zebrafish and is depicted on top of the individual example’s three-dimensional reconstruction. (G) The ratio of silent neurons in relation to their normalized medio-lateral location along both habenula hemispheres of 3,6 and 21dpf. Shading represent SEM. Averages of the two most medial and two most lateral location bins were compared within one age. *p<0.05 for 3dpf, **p<0.01 for 21dpf, Wilcoxon signed-rank test. (H) Confocal microscopy projections of habenula in Tg(vglut2a:dsRed) and Tg(Her4:GFP) double transgenic zebrafish at 3,6 and 21dpf, displaying the glutamatergic neurons (magenta) and potential neuronal progenitors of habenula labelled by Her4 expression (cyan), dorsal view. Bar represents 50μm. White dashed lines delineate the borders of habenula. Note the medial position of neural progenitors, close to the location of silent habenular neurons in (F). (I) Schematic representation of BrdU pulse labelling protocol at 3, 5, 7, 14 and 20dpf. Animals were raised to 21dpf before imaging. (J) Confocal microscopy projections of 21 dpf habenula (white dashed line) showing distribution of neurons that are born at different developmental stages. Birthdating of neurons are achieved by using BrDU (red) pulse labelling. DAPI label marks all cells in blue. Developmental stages, when BrdU was applied, are indicated in each panel. Dorsal view. Bar represents 50μm. (K) Schematic representation of the sequential arrangement of neurons that are born at different developmental stages along the medio-lateral axis, dorsal view. All data presented as mean ±SEM.

Since the older zebrafish habenula exhibits shorter bursts of spontaneous activity with higher frequencies, we asked whether such temporal features of spontaneous activity can also be predictive for the age of habenular neurons. We hypothesized that those habenular neurons with high frequency and short duration burst, might represent the most mature neurons that are born at early development. Whereas neurons with slow and low frequency bursts of activity might represent younger neurons that are born only at later developmental stages. We found that the habenular neurons with distinct temporal features of spontaneous activity were spatially organized at all developmental stages (Figure 5F). This spatial distribution was quantified by using the measure of focality index, ranging from 0 (for random spatial distribution) to 1 (for an extremely focal point) (Yaksi et al., 2007). At all developmental stages the temporal features of habenular spontaneous activity bursts, frequency and duration, were highly focal and not randomly distributed (Figure 5F, mean focality values). Interestingly we also observed that the silent habenular neurons (black dots in Figure 5F), were located closer to the medial wall of habenula (Figure 5F,G), near habenular neural progenitors labelled by Tg(her4.1:GFP) (Yeo et al., 2007) expression (Figure 5H). These results suggest that the changing temporal and spatial features of spontaneous habenular activity can indeed be related to the developmental time point, when the individual habenular neurons are born. To investigate the relationship between the birthdate of habenular neurons and their spatial distribution, we labelled habenular neurons born during different developmental stages using BrdU incorporation (Figure 5I). BrdU birthdating showed that the habenular neurons that are born during distinct developmental stages were spatially clustered (Figure 5J). Moreover, we observed that the birthdate of habenular neurons is a good indicator for their spatial location, where oldest neurons (born early in development) occupied the most lateral wall of habenula, and youngest neurons (born late in development) were closest to the medial wall (Figure 5K). Altogether, our results revealed that the developmental changes in the temporal features of spontaneous habenular activity are spatially organized, and these features could be related to the specific birthdate of habenular neurons at different developmental stages.

### Habenular neurons born at a distinct developmental stage, form a distinct functional cluster with correlated spontaneous activity and similar functional features

Our results above suggested a direct relationship between the birthdate of habenular neurons and their spatial location and functional properties. Based on these results, we hypothesized that the habenular neurons that are born together would also form distinct functional clusters in habenula, with strongly synchronized spontaneous activity. In order to test this, we adopted a previously described method to label cohorts of mammalian cortical neurons born at distinct developmental stages in living animals (Govindan et al., 2018). To label neurons that are born at an early developmental stage, we injected a fluorescent far-red cell tracer in the telencephalic ventricle of 2,5dpf zebrafish larvae (Olstad et al., 2019), near the neural progenitor zone on the medial wall of habenula (Figure 6A). 12 hrs post injection, only the cells near the medial wall of habenula, were labelled with the cell tracer (Figure 6B). We raised these cell tracer injected zebrafish larvae until 21 dpf, and visualized the spatial location of these birthdated neurons in living juvenile zebrafish. In line with our BrdU labelling (Figure 5I,J), we observed that these neurons born at 2,5-3 days were mostly located near the lateral wall of habenula (Supplementary figure 2). Next, we record the spontaneous activity of all juvenile zebrafish habenular neurons expressing GCaMP6s in green spectrum, while identifying their birthdate in far-red spectrum (Figure 6C). We observed striking differences in the functional features of the spontaneous activity between early born (tagged) and late born (untagged) habenular neurons. Significantly larger portion of early born habenular neurons exhibited stronger spontaneous activity(Figure 6D-E) with higher frequency and duration (Figure 6F-G). Moreover, early born habenular neurons also exhibit significantly stronger pairwise correlation, when compared to the rest of habenular neurons (Figure 6H).

These results supported the idea that distinct functional clusters of habenular neurons are born at different developmental stages. To directly test this, we asked whether the developmentally tagged neurons are also related to the spatially organized functional clusters within habenula (Jetti et al., 2014) (Figure 4A-D). To this end, we clustered the spontaneous habenular activity using k-means clustering (Figure 6I-K) and calculated the overlap between developmentally tagged neurons and functional clusters, using a ‘cluster selectivity’ index (Figure 6L). If all developmentally tagged neurons are in only one functional cluster, “clusters selectivity” would result in 1, a random distribution into all clusters would result in 0. We observed that the developmentally tagged cells overlap with significantly fewer clusters compared to the untagged group (Figure 6I-L). These results show for the first time that distinct functional clusters of spatially organized habenular neurons are born at a distinct developmental stage, and exhibit similar features of spontaneous activity. Accumulating evidence suggest that the distinct functional clusters of habenular neurons mediate distinct behaviours (Agetsuma et al., 2010; Amo et al., 2014; Andalman et al., 2019; Chou et al., 2016; Duboue et al., 2017; Lee et al., 2010). Sequential addition of new functional clusters with new properties across development could therefore explain the expansion of zebrafish behavioural repertoire across development (Dreosti et al., 2015; Valente et al., 2012).

**Figure 6:**
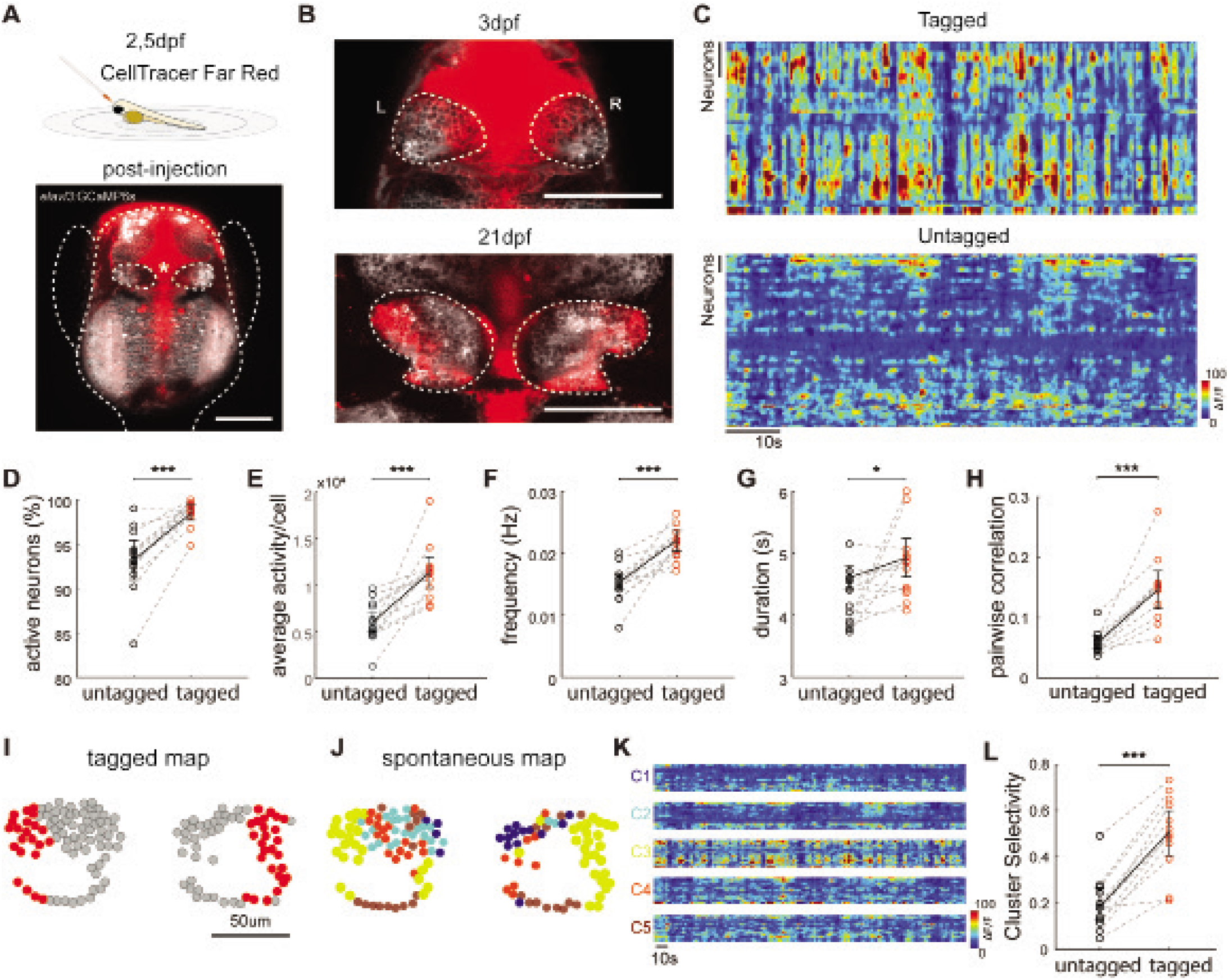
Habenular neurons that are born at a distinct developmental stage exhibit similar functional properties and more correlated activity. (A) Top panel shows a schematic representation of cell Tracer injections at 2,5dpf zebrafish. Bottom panel shows confocal microscopy image of Tg(elavl3:GCaMP6s) zebrafish labelling neurons (white) displaying the ventricle that was injected with the cell tracer (red), dorsal view. Asterix shows injection site in the ventricle near habenula that is delineated by dashed white line. Bar represents 100μm. Borders of the zebrafish are delineated by a dashed white line. (B) Confocal microscopy images of zebrafish habenula at 3 (top) and 21 (bottom) dpf, after the injection of the cell tracer at 2,5 dpf. Dorsal view. Habenula is delineated by a dashed white line. Bars represent 100μm. L,left; R;right. (C) Spontaneous activity of tagged habenular neurons in 21dpf zebrafish that were birthdated by the cell tracer injection at 2,5 dpf (top) and the remaining untagged habenular neurons (bottom). Bars represent 10 neurons. Spontaneous neural activity levels are colour coded for each neuron. (D) The ratio of spontaneously active neurons with at least one significant spontaneous activity burst in the 21 dpf zebrafish habenula. Note that a smaller ratio of untagged late-born neurons (black) were active when compared to tagged early born (red) neurons by the celltracer injection at 2,5 dpf (n=12 fish). (E) Average spontaneous activity that is represented as total area under the curve of all detected events, in habenular neurons that are developmentally tagged versus untagged. (F) Average frequency of spontaneous activity bursts in 21dpf habenular neurons that are developmentally tagged versus untagged. (G) Average duration of spontaneous activity bursts in 21dpf habenular neurons that are developmentally tagged versus untagged.(H) Average pairwise correlations of tagged versus untagged habenular neurons in 21dpf zebrafish.*p<0.05, ***p<0.001, Wilcoxon signed-rank test. (I) Spatial distribution of tagged versus untagged habenular neurons, in an example of a 21 dpf zebrafish. (J) Spatial distribution of habenular neurons colour coded according to their functional clusters based on their spontaneous activity, using k-means clustering in the same example zebrafish as in I. (K) Colour coded neural traces showing the spontaneous activity of each habenular neuron that belong to a given cluster “C”, in the same example zebrafish as I and J. Warm colours represents increased neural activity. Note the prominent overlap between the functional cluster C3 (yellow neurons in J) and the cell tracer tagged neurons in I. (L) Cluster selectivity, depicting how tagged neurons (red) are distributed into functional clusters based on their spontaneous activity, when compared to untagged neurons. High selectivity means that tagged neurons belong to fewer clusters, when compared to untagged neurons (black dots), ***p<0.001, Wilcoxon signed-rank test. All data presented as mean ±SEM.

## DISCUSSION

In this study we investigated the functional development of habenular circuits across multiple developmental stages of zebrafish from relatively simpler larvae to juvenile zebrafish with complex behaviours (Amo et al., 2014; Andalman et al., 2019; Chou et al., 2016; Dreosti et al., 2015; Hinz and de Polavieja, 2017; Ksenia Yashina, 2019; Valente et al., 2012). The use of juvenile zebrafish is getting increasingly popular due to their transparent brains and expanded behavioural repertoire that requires habenular function (Andalman et al., 2019; Dreosti et al., 2015; Hinz and de Polavieja, 2017; Valente et al., 2012; Yashina et al., 2019). Our results revealed that as the zebrafish develop from larval to juvenile stage, habenular circuits undergo multiple transitions in its architecture, sensory computations and intrinsically generated spontaneous activity, which could support the expansion of the behavioural repertoire in developing zebrafish.

Previous studies indicated the presence of olfactory (Dreosti et al., 2014; Jetti et al., 2014; Krishnan et al., 2014) and visual (Cheng et al., 2017; Dreosti et al., 2014; Zhang et al., 2017) responses in habenula, which were suggested to be involved in generating odour avoidance (Krishnan et al., 2014), light preference (Zhang et al., 2017) and circadian behaviours (Lin and Jesuthasan, 2017). While these critical sensory-evoked behaviours are present already in larval zebrafish, the nature of sensory computations and how such computations maturate across development have not been investigated. Our results revealed that at early developmental stages habenula is mostly innervated by the visual inputs, hence responding to visual stimuli. However, as the olfactory inputs and responses become prominent across development, habenula transitions into a multi-sensory processing brain region, which can integrate both visual and olfactory stimuli. Interestingly, already from the early developmental stages, we observe that visual and olfactory information are distinctly encoded by the activation of separate neural populations located in different habenular hemispheres. Such distinct encoding of different sensory modalities at all developmental stages, highlights the important role of habenula in olfactory and visually-driven behaviours (Cheng et al., 2017; Dreosti et al., 2014; Jetti et al., 2014; Krishnan et al., 2014; Zhang et al., 2017).

As in other lower vertebrates(Concha and Wilson, 2001), zebrafish dorsal habenula exhibits prominent molecular (Aizawa et al., 2012; deCarvalho et al., 2014; Pandey et al., 2018; Viswanath et al., 2013) and structural asymmetries (Bianco et al., 2008). Not surprisingly, habenular lateralization has important consequences with respect to the segregation of different sensory modalities across habenular hemispheres and for regulating the targeting of habenular axons to different output regions (Amo et al., 2014; Bianco et al., 2008; Duboue et al., 2017). We observed that the functional lateralization of sensory inputs and computations in habenula is prominent mostly in the dHb, which develops early during development (Aizawa et al., 2007). Whereas vHb, which was suggested to develop later (Amo et al., 2010; Beretta et al., 2013), exhibits no such functional lateralization, despite the prominent sensory evoked responses. This developmental order eventually leads to a decrease in functional lateralization across the entire habenula as the animals maturate. Interestingly, like in sensory systems (Moorman et al., 1999; Webster and Webster, 1977; Wiesel and Hubel, 1965), maturation of zebrafish habenula requires the early presence of sensory inputs (Dreosti et al., 2014). When these inputs are removed before a critical time window, zebrafish dHb loses it functional lateralization (Dreosti et al., 2014). Our observations are also in line with the lack of evidence for the functional lateralization in the mammalian habenula, where most studies focus on the lateral habenula (Huang et al., 2019). Hence, the ontogeny of zebrafish habenular function might follow the phylogeny of vertebrate habenula evolution. We argue that the apparent lack of lateralization in mammalian habenula is due to continued maturation of lateral habenula and reduced availability of sensory information during mammalian in utero development.

Activity-dependent maturation of neural circuits, do not solely rely on sensory inputs but also include endogenously generated spontaneous activity (Penn and Shatz, 1999). In several brain regions spontaneously generated activity is thought to play important roles in the maturation of synaptic connections, refining network topography (Katz and Shatz, 1996; Moreno-Juan et al., 2017; Xu et al., 2011) and entraining developing circuits (Clause et al., 2014; Gretenkord et al., 2019; Huberman et al., 2006). In our study, we observed that habenular networks exhibit spatially organized spontaneous activity already at early developmental stages. In fact, a recent transcriptomics study revealed a topographic organization of habenular neurons based on their distinct molecular features already at 10 days old zebrafish larvae (Pandey et al., 2018), which likely overlap with the topographically organized spontaneous activity we observed here. Interestingly we also observed that temporal features of habenular spontaneous activity changed, leading to faster kinetics at older ages. The trend for faster kinetics of intrinsically generated activity across development was also observed in other higher brain regions, such as the hippocampus (Buzsaki and Draguhn, 2004; Leinekugel et al., 1998). What may underlie the faster kinetics of spontaneous activity bursts in older ages ? It is likely that multiple processes can modulate spontaneous habenular activity leading to the observed dynamical changes during development. One possibility is that early spontaneous activity might be driven by glia (Zhang et al., 2019) or other support cells (Babola et al., 2018), which usually generate slow bursts of activity. In fact, it was previously shown that astrocytes can modulate the bursting activity in the rodent habenula (Cui et al., 2018) and astroglia activation in zebrafish can excite nearby neurons (Verdugo et al., 2019). Alternatively, it is possible that as the habenula develops, synaptic connections between habenular neurons become more mature. Together with the increased inhibition we observed in the habenula, by the addition of GABAergic connections, such mature synapses can support faster neural dynamics. Finally, it is likely that part of the habenular spontaneous activity is driven by the activation of the ancestral cortico-limbic homologs in zebrafish forebrain. For example, it was shown that habenula is strongly driven by inputs received from entopeduncular nucleus (Herkenham and Nauta, 1977; Hong and Hikosaka, 2008; Turner et al., 2016) lateral hypothalamus (Lazaridis et al., 2019) and cortex (Warden et al., 2012). Hence it is likely that the homologues of these input regions are not yet fully developed and cannot drive habenular activity in young zebrafish larvae. A recent study showed that the neurons in the zebrafish equivalent of amygdala (Dm), which drives both entopeduncular nucleus and lateral hypothalamus (Lal et al., 2018), also develop during late juvenile stage. Taken together, it is very likely that all these processes jointly contribute to the habenular spontaneous activity. How each of these above mentioned factors differentially affect the spontaneous activity and maturation of habenular circuits is yet to be investigated in future studies.

Our results revealed that the developmental changes in the temporal kinetics of habenular spontaneous activity is accompanied by a sequential addition of new born neurons across development. More specifically, we found that the spatially confined locations of neuronal clusters that are born during distinct developmental stages, follow a medio-lateral arrangement. In this arrangement youngest neurons are located closer to the neural progenitors of the habenula on the medial wall, while the oldest habenular neurons are located closer to lateral wall. Similar sequential stacking of newly born neurons was also observed during forebrain development in both zebrafish (Furlan et al., 2017) and in the cortex of rodents (Angevine and Sidman, 1961; Lein et al., 2017; Luskin and Shatz, 1985). Moreover, such sequential order of neurogenesis contributes to different neural subtypes in the rodent entorhinal cortex (Donato et al., 2017). Hence it is possible that recently described populations of habenular neurons with distinct molecular identities (Pandey et al., 2018) might be born during different developmental stages of the zebrafish habenula. Topographically organized molecular (Pandey et al., 2018) and functional clusters (Andalman et al., 2019; Jetti et al., 2014; Pandey et al., 2018) of habenular neurons represent distinct subdomains or functional modules within habenula, and are most likely associated with different aspects of animal behaviour. Our data revealed that some of the functional clusters within habenula are born at a distinct developmental stage, stay together in a spatially confined location and exhibit highly synchronized spontaneous activity. We therefore propose that distinct functional clusters of habenular neurons with specific features of activity are preferentially born at different developmental time points. In turn, such a developmental order would contribute to an increased diversity of habenular neurons across development. Several recent studies suggest that complex and cognitively demanding behaviours arise later in development as animals maturate (Amo et al., 2014; Andalman et al., 2019; Valente et al., 2012; Yashina et al., 2019). Our findings in habenula suggest that such expansion of animal behaviour might be due to incorporation of new functional modules at different developmental time points. In the future, it will be interesting to see whether such sequential addition of new functional modules with distinct roles in regulating animal behaviour is a feature that is not only unique to habenula but is preserved across the brain.

## ACKNOWLEDGEMENTS

We thank M. Ahrens (HHMI, Janelia Farm, USA), C. Wyart (ICM, Paris, France), H. Baier (MPI, Martinsried, Germany) and Shin-ichi Higashijima (Okazaki Institute for Integrative Bioscience, Japan) for transgenic lines. We thank S. Eggen, M. Andresen, V. Nguyen and our fish facility support team for technical assistance. We thank Nathalie Jurisch-Yaksi for feedback on the manuscript. We thank the Yaksi lab members for stimulating discussions. This work was funded by ERC starting grant 335561 (S.F., E.Y.). Work in the E.Y. lab is funded by the Kavli Institute for Systems Neuroscience at NTNU.

## AUTHOR CONTRIBUTIONS

Conceptualization, S.F., E.Y.; Methodology and data, S.F., M.I.C. C.K.; Data Analysis, S.F., C.D.V.; Investigation, all authors; Writing, S.F., E.Y.; Review & Editing, all authors; Funding Acquisition and Supervision, E.Y.

## DECLARATION OF INTERESTS

The authors declare no competing interests.

## STAR METHODS

### Contact for Reagent and Resource Sharing

Further information and requests for reagents may be directed to, and will be fulfilled by the lead author Emre Yaksi.

### Fish maintenance

The animal facilities and maintenance of zebrafish, Danio rerio, were approved by the Norwegian Food Safety Authority. Fish were kept in 3.5liter tanks at a density of 15-20 fish per tank in a Techniplast Zebtech Multilinking system at 28°C, pH 7, 6.0ppm O2 and 700*μ*S, at a 14:10 hour light/dark cycle. Fish received a normal diet of dry food (Zebrafeed, Sparos) two times per day and were after 6dpf also fed with Artemia nauplii once a day (Grade0, platinum Label, Argent Laboratories). Larvae were maintained in egg water (1.2g marine salt and 0.1% methylene blue in 20L RO water) from fertilization to 3dpf. From 3dpf to 6dpf larvae were kept in AFW (1.2g marine salt in 20L RO water). Animals used for experiments at 21dpf were transferred to 3.5 liter tanks in the zebrafish facility at 3dpf.

All experimental procedures performed on zebrafish larvae and juveniles were in accordance with the directive 2010/63/EU of the European Parliament and the Council of the European Union and approved by the Norwegian Food Safety Authorities and Landesdirektion Sachsen, Germany (permit numbers TVV-52/2015 and TVV-35/2016).

For experiments the following fish lines were used: Tg(elavl3:GCaMP6s) (Vladimirov et al., 2014), Tg(gad1b:GFP) (Satou et al., 2013), Tg(vglut2a:dsRed) (Miyasaka et al., 2009), Tg(Lhx2a:gap-YFP) (Miyasaka et al., 2009), Et(−0.6hsp70l:Gal4-VP16) s1020t; UAS:nfsB-mCherry (Scott and Baier, 2009), Tg(her4.1:GFP) (Yeo et al., 2007). Experiments were performed on embryos of nacre (mitfa(Lister et al., 1999)) background.

### Confocal anatomical imaging

Prior to embedding, fish were anaesthetized with 0.02% MS222. Animals were then embedded in 1% (for3-6dpf) or 2% (for 21dpf) low-melting-point agarose (LMP, Fisher Scientific) in a recording chamber (Fluorodish, World Precision Instruments) with AFW. Anatomical Z-scans were acquired using a Zeiss Examiner Z1 confocal microscope with a 20x water-immersion objective (Zeiss, NA 1.0, Plan-Apochromat) at room temperature, using 4-10x average for each plane.

### BrDU stainings and imaging

Labelling of newborn neurons with BrdU was performed at 3,5; 5,5; 7,5; 14,5; 20,5dpf in Tg(HuC:GFP) (Park et al., 2000) fish outcrossed to wild type (AB) fish. 25-hpf embryos were dechorinated by Proteinase K (0.1 mg/ml) (Cat#:25530049) at room temperature and washed several times with 1x E3 medium. A stock of 10 mM BrdU (Sigma, Cat# B5002) in 1x E3 medium was prepared, aliquoted, and stored at −20 °C until use. For each treatment, a final concentration of 500 μM BrdU was applied to 3, and 5 dpf fish. For 7, 14 and 21 dpf fish, 2.5 mM BrdU was used. Treatments were all performed for 5 hours. Fish were kept in a dark incubator until 5 dpf and were transferred to the water system at 6 dpf. From this point on, 14/10 hours light/dark cycle was maintained. At 22 dpf, fish were sacrificed and fixed in 4% PFA-1% DMSO at +4 °C overnight, washed with 0.8% Triton-X in 1X PBS (PBSTx), dehydrated by a serial methanol gradient, and kept at −20 °C until further use. For immunohistochemical stainings, fish were rehydrated back to aqueous phase (1X PBS). Brains were dissected out in cold 1x PBS, and were incubated in pre-cooled acetone for 10 mins at −20°C. Brains were incubated in preheated 2mM HCl at 37 °C for 12 minutes, were cooled at room temperature for 10 min, and washed with 0.8% PBSTx. Samples were incubated in Rat anti-BrdU IgG (1:200 Bio-Rad, Cat# MCA2060) monoclonal antibody overnight at +4 °C. After serial washes in 0.8% PBSTx, the secondary antibody (Goat anti-rat Alexa 555, 1:500 dilution, ThermoFishcher, Cat#A-21434) and DAPI at 1:3000 dilution were applied. Following an overnight incubation at +4 °C, samples were washed for a total of 2 hours with 0.8% PBSTx. For preservation, samples were transferred to 80% Glycerol in 1X PBS and were kept at +4 °C in the dark. For imaging, labelled brains were mounted in 80% glycerol on a glass slide with cover slip. Anatomical Z-scans of habenula were acquired using a Zeiss Examiner Z1 confocal microscope with a 20x plan NA 0.8 objective, using 4-10x average for each plane.

### In vivo two-photon calcium imaging and sensory stimulation

Two-photon calcium imaging was performed on 3dpf, 6dpf, and 21dpf old Tg(elavl3:GCaMP6s) zebrafish. Prior to embedding, fish were anaesthetized with cold AFW. Animals were then embedded in 1% (for 3-6dpf) and 2% (for 21dpf) low-melting-point agarose (LMP, Fisher Scientific) in a recording chamber (Fluorodish, World Precision Instruments). Low-melting-point agarose solidified for 20min, and the section covering the nose was carefully removed to expose the nostrils. The animal was then placed under the microscope with constant perfusion of AFW bubbled with carbogen (95% O2 and 5%CO2). First spontaneous activity was measured for 30min. Afterwards, 5 repetitions of sensory stimuli (red light flash or food odour) were applied. The food odour stimulus was prepared by adding 1g of standard dried fish food (Zebrafeed, Sparos, <100), in 50ml of AFW, dissolved for 1hr and filtered with a 22μm filter. This was then further diluted 1:50 in AFW. The odour stimulus was delivered for 5sec through a tube positioned in the front of the fish, connected to an Arduino Due-controlled HPLC injection valve (Valco Instruments). Fluorescein (10-4M) was dissolved in AFW and used to measure precise onset of odour delivery to the nose at the end of each experiment. For the light stimulus we used a red LED (LZ1-00R105, LedEngin; 625 nm wavelength) and placed in the front of the recording chamber near the tube. The light stimulus was a square-wave flash of 2s duration with an intensity of 450mW. The recordings were performed with a two-photon microscope (Scientifica) using a 16x water immersion objective (Nikon, NA 0.8, LWD 3.6,plan). A mode-locked Ti:Sapphire laser (MaiTai Spectra-Physics) tuned to 920nm was used for excitation. Volumetric recordings (8 planes with Piezo) were obtained at an acquisition rate of 31.9Hz for a volume of 1536×512 pixels x 8 planes. Total duration of the recordings was 45-60min.

### In vivo birthdating of neurons

Prior to injections, 2.5dpf larvae were dechorionated manually and anaesthetized in 0.01% MS-222 in AFW for 5-10min. Injections were done on larvae embedded in 1% low-melting point agarose in AFW and 0.01% MS-222 in AFW. The stock solution contained 5mM CellTrace Far Red(Thermofisher) dissolved in DMSO provided with the proliferation kit. The injection mixtures contained 0.5μl of this CellTrace Far Red/DMSO stock solution dissolved in 2.5μl of artificial cerebrospinal fluid (aCSF) with a final concentration of 1mM. The aCSF contained the following: 124 mM NaCl, 22 mM D-(+)-Glucose, 2.0 mM KCl, 1.6 mM MgSO4 • 7 H2O, 1.3 mM KH2PO4, 24 mM NaHCO3, 2.0 mM CaCl2 ‣ 2 H2O. The injection needles were pulled with a Sutter Instrument Co. Model P-2000, from thin-walled borosilicate capillaries (1.00 mm; VWR), with the following settings: heat = 785, filament = 4, velocity = 40, delay = 220, pull = 70. The needle tip was cut open with forceps afterwards and a pressure injector (Eppendorf Femtojet 4i) was used to inject 1 nL of solution in the telencephalic ventricle near habenula (Olstad et al., 2019). The pressure and time used for the injection were calibrated for each needle using a 0.01 mm calibration slide for microscopy. Usually, the pressure ranged between 100-150 hPa and the time span of the pressure pulse lasted for 0.30 – 0.70 s. After injection, fish were released from agarose and recovered at 28°C AFW and transferred to the fish facility.

From 19-25dpf fish were prepared for imaging as previously described for calcium imaging. Imaging was performed using a Zeiss Examiner Z1 confocal microscope with a 20x water-immersion objective (Zeiss, NA 1.0, Plan-Apochromat) at room temperature and constant perfusion of AFW bubbled with carbogen (95% O2 and 5% CO2). In vivo calcium recordings of the spontaneous activity in habenula were acquired at an acquisition rate of 1.918Hz for 512×1024pixels during 5min.

### Data Analysis

Two-photon microscopy images were aligned using an adapted algorithm (Reiten et al., 2017) that corrects for occasional drift in the XY dimension, based on “hierarchical model-based motion estimation (Bergen et al., 1992). Every recording was manually checked for motion and corresponding frames were discarded from further analysis. Individual neurons were semi-automatically detected using a pattern recognition as previously described (Jetti et al., 2014; Reiten et al., 2017; Verdugo et al., 2019). Once neurons were detected, their locations were individually tracked during the recording. If any z-motion drift or if some detected neurons were no longer visible those neurons were discarded from the further analysis. The pixels belonging to each neuron were then averaged providing the complete time course of each individual neuron over time. Same cell-detection algorithms were used for the confocal recordings in Figure 6. To distinguish between developmentally tagged and untagged neurons, we manually identified the red cells. To normalize the spontaneous activity for each cell to its baseline fluorescence, we calculated the fractional change in fluorescence (ΔF/F) relative to the baseline, which was calculated as the 8th percentile of activity observed within 2min time window as preciously described (Romano et al., 2017). Clustering of neurons was performed as previously reported based on the k-means clustering algorithm in MATLAB as well as the calculation of cluster fidelity and cluster selectivity for sensory and tagged cells (Jetti et al., 2014).

Prior to event detection, traces from neurons in the two-photon recordings were resampled to a final rate of 2 Hz (using decimate function in MATLAB), to match the sampling rate with confocal experiments. Significant calcium events were detected using an algorithm (Romano et al., 2017) that detects calcium events significantly different from noise level within a 95% confidence interval. A cell was considered active if at least one event was detected in 4min of ongoing activity. The amplitude of events was defined as the maximum peak in the event. The average activity per cell was defined as the sum of the area under the curve (using trapezoidal numerical integration method: function ‘trapz’ in MATLAB) for all events within one neuron. The fractional change in fluorescence (ΔF/F) for the odour and light stimulus was calculated by subtracting the average baseline fluorescence before stimulus onset (5sec) from the responsive window (10 sec for odour and 2sec for light from the onset of the stimulus). Cells were classified as responding if at least 4 out of 5 trials showed the same sign difference (+/-) locked to the stimulus onset. Only the positively responding neurons were taken into account for further analysis.

The focality index was calculated as 1 minus the average of the Euclidean distances between the top 10% neurons (with highest frequencies or durations) divided by the average of the Euclidean distances of all neurons within each hemisphere. The average focality index per fish was calculated by taking the average over the two hemispheres. Same focality index was calculated for the sensory-responding cells.

Cluster fidelity was calculated by measuring the probability of pairs of neurons being in the same cluster during two different time periods (Jetti et al., 2014). As a control, we shuffled the cluster identity of neurons randomly.

To calculate cluster selectivity, we first calculated the cluster identity of each neuron using k-means clustering of spontaneous neural activity(Jetti et al., 2014). Later we identified neurons with a given property, in this case either sensory responding neurons (Figure 4G), of developmentally tagged neurons (Figure 6L) and measured the sparseness in the distribution of identified neurons into clusters(Jetti et al., 2014). If sparseness is 1 all identified neurons are selectively member of a single cluster, if sparseness is 0 all identified neurons are equally distributed into all functional clusters identified by k-means clustering.

In the confocal microscopy functional imaging, results for frequency, durations, number of active neurons, average activity and pairwise correlations, were averaged over the planes imaged in each fish, as these recordings were done at different timepoints.

The lateralization index was calculated as the difference between the percentage of responding neurons in left and right hemispheres.

### Quantification and Statistical Analysis

Statistical analysis was done using MATLAB and p-values are indicated in the figure legends (*p<0.05, **p<0.01, ***p<0.001). Student t-test was used for paired data and unpaired t-test when data was obtained from two independent datasets. For the data that displayed no Gaussian distribution, we used Wilcoxon signed rank test for paired data and Wilcoxon ranksum test for unpaired data.

### Data and software availability

All analysis was performed with Fiji and MATLAB as indicated in the results section. All custom MATLAB scripts are available upon request.

## Notes

### Competing Interest Statement

The authors have declared no competing interest.

### Summary of Updates

We uploaded the original version of our manuscript in version 1, as it was asked by the scientific journal reviewing our manuscript.

## REFERENCES

Agetsuma, M., Aizawa, H., Aoki, T., Nakayama, R., Takahoko, M., Goto, M., Sassa, T., Amo, R., Shiraki, T., Kawakami, K., et al. (2010). The habenula is crucial for experience-dependent modification of fear responses in zebrafish. Nat Neurosci 13, 1354–1356.

Ahumada-Galleguillos, P., Lemus, C.G., Diaz, E., Osorio-Reich, M., Hartel, S., and Concha, M.L. (2017). Directional asymmetry in the volume of the human habenula. Brain Struct Funct 222, 1087–1092.

Aizawa, H., Goto, M., Sato, T., and Okamoto, H. (2007). Temporally regulated asymmetric neurogenesis causes left-right difference in the zebrafish habenular structures. Dev Cell 12, 87–98.

Aizawa, H., Kobayashi, M., Tanaka, S., Fukai, T., and Okamoto, H. (2012). Molecular characterization of the subnuclei in rat habenula. J Comp Neurol 520, 4051–4066.

Altman, J., Brunner, R.L., and Bayer, S.A. (1973). The hippocampus and behavioral maturation. Behav Biol 8, 557–596.

Amo, R., Aizawa, H., Takahoko, M., Kobayashi, M., Takahashi, R., Aoki, T., and Okamoto, H. (2010). Identification of the zebrafish ventral habenula as a homolog of the mammalian lateral habenula. J Neurosci 30, 1566–1574.

Amo, R., Fredes, F., Kinoshita, M., Aoki, R., Aizawa, H., Agetsuma, M., Aoki, T., Shiraki, T., Kakinuma, H., Matsuda, M., et al. (2014). The habenulo-raphe serotonergic circuit encodes an aversive expectation value essential for adaptive active avoidance of danger. Neuron 84, 1034–1048.

Andalman, A.S., Burns, V.M., Lovett-Barron, M., Broxton, M., Poole, B., Yang, S.J., Grosenick, L., Lerner, T.N., Chen, R., Benster, T., et al. (2019). Neuronal Dynamics Regulating Brain and Behavioral State Transitions. Cell 177, 970–985 e920.

Anderson, J.R. (1984). The development of self-recognition: a review. Dev Psychobiol 17, 35–49.

Angevine, J.B., Jr., and Sidman, R.L. (1961). Autoradiographic study of cell migration during histogenesis of cerebral cortex in the mouse. Nature 192, 766–768.

Babola, T.A., Li, S., Gribizis, A., Lee, B.J., Issa, J.B., Wang, H.C., Crair, M.C., and Bergles, D.E. (2018). Homeostatic Control of Spontaneous Activity in the Developing Auditory System. Neuron 99, 511–524 e515.

Ben-Ari, Y. (2001). Developing networks play a similar melody. Trends Neurosci 24, 353–360.

Beretta, C.A., Dross, N., Bankhead, P., and Carl, M. (2013). The ventral habenulae of zebrafish develop in prosomere 2 dependent on Tcf7l2 function. Neural Dev 8, 19.

Bergen, J.R., Anandan, P., Hanna, K.J., and Hingorani, R. (1992). Hierarchical model-based motion estimation (Berlin, Heidelberg: Springer Berlin Heidelberg).

Bianco, I.H., Carl, M., Russell, C., Clarke, J.D., and Wilson, S.W. (2008). Brain asymmetry is encoded at the level of axon terminal morphology. Neural Dev 3, 9.

Bjerknes, T.L., Dagslott, N.C., Moser, E.I., and Moser, M.B. (2018). Path integration in place cells of developing rats. Proc Natl Acad Sci U S A 115, E1637–E1646.

Brenowitz, E.A., and Beecher, M.D. (2005). Song learning in birds: diversity and plasticity, opportunities and challenges. Trends Neurosci 28, 127–132.

Burgess, H.A., and Granato, M. (2007). Sensorimotor gating in larval zebrafish. J Neurosci 27, 4984–4994.

Buzsaki, G., and Draguhn, A. (2004). Neuronal oscillations in cortical networks. Science 304, 1926–1929.

Cheng, R.K., Krishnan, S., Lin, Q., Kibat, C., and Jesuthasan, S. (2017). Characterization of a thalamic nucleus mediating habenula responses to changes in ambient illumination. BMC Biol 15, 104.

Chou, M.Y., Amo, R., Kinoshita, M., Cherng, B.W., Shimazaki, H., Agetsuma, M., Shiraki, T., Aoki, T., Takahoko, M., Yamazaki, M., et al. (2016). Social conflict resolution regulated by two dorsal habenular subregions in zebrafish. Science 352, 87–90.

Clause, A., Kim, G., Sonntag, M., Weisz, C.J., Vetter, D.E., Rubsamen, R., and Kandler, K. (2014). The precise temporal pattern of prehearing spontaneous activity is necessary for tonotopic map refinement. Neuron 82, 822–835.

Concha, M.L., and Wilson, S.W. (2001). Asymmetry in the epithalamus of vertebrates. J Anat 199, 63–84.

Cui, Y., Yang, Y., Ni, Z., Dong, Y., Cai, G., Foncelle, A., Ma, S., Sang, K., Tang, S., Li, Y., et al. (2018). Astroglial Kir4.1 in the lateral habenula drives neuronal bursts in depression. Nature 554, 323–327.

deCarvalho, T.N., Subedi, A., Rock, J., Harfe, B.D., Thisse, C., Thisse, B., Halpern, M.E., and Hong, E. (2014). Neurotransmitter map of the asymmetric dorsal habenular nuclei of zebrafish. Genesis 52, 636–655.

Donato, F., Jacobsen, R.I., Moser, M.B., and Moser, E.I. (2017). Stellate cells drive maturation of the entorhinal-hippocampal circuit. Science 355.

Dreosti, E., Lopes, G., Kampff, A.R., and Wilson, S.W. (2015). Development of social behavior in young zebrafish. Front Neural Circuits 9, 39.

Dreosti, E., Vendrell Llopis, N., Carl, M., Yaksi, E., and Wilson, S.W. (2014). Left-right asymmetry is required for the habenulae to respond to both visual and olfactory stimuli. Curr Biol 24, 440–445.

Duboue, E.R., Hong, E., Eldred, K.C., and Halpern, M.E. (2017). Left Habenular Activity Attenuates Fear Responses in Larval Zebrafish. Curr Biol 27, 2154–2162 e2153.

Elstrott, J., Anishchenko, A., Greschner, M., Sher, A., Litke, A.M., Chichilnisky, E.J., and Feller, M.B. (2008). Direction-selectivity in the retina is established independent of visual experience and cholinergic retinal waves. Neuron 58, 499–506.

Feller, M.B. (1999). Spontaneous correlated activity in developing neural circuits. Neuron 22, 653–656.

Flanigan, M., Aleyasin, H., Takahashi, A., Golden, S.A., and Russo, S.J. (2017). An emerging role for the lateral habenula in aggressive behavior. Pharmacol Biochem Behav 162, 79–86.

Furlan, G., Cuccioli, V., Vuillemin, N., Dirian, L., Muntasell, A.J., Coolen, M., Dray, N., Bedu, S., Houart, C., Beaurepaire, E., et al. (2017). Life-Long Neurogenic Activity of Individual Neural Stem Cells and Continuous Growth Establish an Outside-In Architecture in the Teleost Pallium. Curr Biol 27, 3288–3301 e3283.

Galli, L., and Maffei, L. (1988). Spontaneous impulse activity of rat retinal ganglion cells in prenatal life. Science 242, 90–91.

Govindan, S., Oberst, P., and Jabaudon, D. (2018). In vivo pulse labeling of isochronic cohorts of cells in the central nervous system using FlashTag. Nat Protoc 13, 2297–2311.

Gretenkord, S., Kostka, J.K., Hartung, H., Watznauer, K., Fleck, D., Minier-Toribio, A., Spehr, M., and Hanganu-Opatz, I.L. (2019). Coordinated electrical activity in the olfactory bulb gates the oscillatory entrainment of entorhinal networks in neonatal mice. PLoS Biol 17, e2006994.

Heap, L.A.L., Vanwalleghem, G., Thompson, A.W., Favre-Bulle, I.A., and Scott, E.K. (2018). Luminance Changes Drive Directional Startle through a Thalamic Pathway. Neuron 99, 293–301 e294.

Herkenham, M., and Nauta, W.J. (1977). Afferent connections of the habenular nuclei in the rat. A horseradish peroxidase study, with a note on the fiber-of-passage problem. J Comp Neurol 173, 123–146.

Hikosaka, O. (2010). The habenula: from stress evasion to value-based decision-making. Nat Rev Neurosci 11, 503–513.

Hinz, R.C., and de Polavieja, G.G. (2017). Ontogeny of collective behavior reveals a simple attraction rule. Proc Natl Acad Sci U S A 114, 2295–2300.

Hong, S., and Hikosaka, O. (2008). The globus pallidus sends reward-related signals to the lateral habenula. Neuron 60, 720–729.

Hong, S., Jhou, T.C., Smith, M., Saleem, K.S., and Hikosaka, O. (2011). Negative reward signals from the lateral habenula to dopamine neurons are mediated by rostromedial tegmental nucleus in primates. J Neurosci 31, 11457–11471.

Huang, L., Xi, Y., Peng, Y., Yang, Y., Huang, X., Fu, Y., Tao, Q., Xiao, J., Yuan, T., An, K., et al. (2019). A Visual Circuit Related to Habenula Underlies the Antidepressive Effects of Light Therapy. Neuron.

Huberman, A.D., Speer, C.M., and Chapman, B. (2006). Spontaneous retinal activity mediates development of ocular dominance columns and binocular receptive fields in v1. Neuron 52, 247–254.

Jabaudon, D. (2017). Fate and freedom in developing neocortical circuits. Nat Commun 8, 16042.

Jetti, S.K., Vendrell-Llopis, N., and Yaksi, E. (2014). Spontaneous activity governs olfactory representations in spatially organized habenular microcircuits. Curr Biol 24, 434–439.

Katz, L.C., and Shatz, C.J. (1996). Synaptic activity and the construction of cortical circuits. Science 274, 1133–1138.

Kerschensteiner, D. (2014). Spontaneous Network Activity and Synaptic Development. Neuroscientist 20, 272–290.

Krishnan, S., Mathuru, A.S., Kibat, C., Rahman, M., Lupton, C.E., Stewart, J., Claridge-Chang, A., Yen, S.C., and Jesuthasan, S. (2014). The right dorsal habenula limits attraction to an odor in zebrafish. Curr Biol 24, 1167–1175.

Ksenia Yashina, Á.T.-C., Andreas Herz, Herwig Baier (2019). Zebrafish exploit visual cues and geometric relationships to form a spatial memory. ISCIENCE.

Lal, P., Tanabe, H., Suster, M.L., Ailani, D., Kotani, Y., Muto, A., Itoh, M., Iwasaki, M., Wada, H., Yaksi, E., et al. (2018). Identification of a neuronal population in the telencephalon essential for fear conditioning in zebrafish. BMC Biol 16, 45.

Lazaridis, I., Tzortzi, O., Weglage, M., Martin, A., Xuan, Y., Parent, M., Johansson, Y., Fuzik, J., Furth, D., Fenno, L.E., et al. (2019). A hypothalamus-habenula circuit controls aversion. Mol Psychiatry.

Lee, A., Mathuru, A.S., Teh, C., Kibat, C., Korzh, V., Penney, T.B., and Jesuthasan, S. (2010). The habenula prevents helpless behavior in larval zebrafish. Curr Biol 20, 2211–2216.

Lein, E.S., Belgard, T.G., Hawrylycz, M., and Molnar, Z. (2017). Transcriptomic Perspectives on Neocortical Structure, Development, Evolution, and Disease. Annu Rev Neurosci 40x, 629–652.

Leinekugel, X., Khalilov, I., Ben-Ari, Y., and Khazipov, R. (1998). Giant depolarizing potentials: the septal pole of the hippocampus paces the activity of the developing intact septohippocampal complex in vitro. J Neurosci 18, 6349–6357.

Lin, Q., and Jesuthasan, S. (2017). Masking of a circadian behavior in larval zebrafish involves the thalamo-habenula pathway. Sci Rep 7, 4104.

Lindsay, S.M., and Vogt, R.G. (2004). Behavioral responses of newly hatched zebrafish (Danio rerio) to amino acid chemostimulants. Chem Senses 29, 93–100.

Lister, J.A., Robertson, C.P., Lepage, T., Johnson, S.L., and Raible, D.W. (1999). nacre encodes a zebrafish microphthalmia-related protein that regulates neural-crest-derived pigment cell fate. Development 126, 3757–3767.

Luskin, M.B., and Shatz, C.J. (1985). Neurogenesis of the cat’s primary visual cortex. J Comp Neurol 242, 611–631.

Maffei, L., and Galli-Resta, L. (1990). Correlation in the discharges of neighboring rat retinal ganglion cells during prenatal life. Proc Natl Acad Sci U S A 87, 2861–2864.

Matsumoto, M., and Hikosaka, O. (2007). Lateral habenula as a source of negative reward signals in dopamine neurons. Nature 447, 1111–1115.

Meister, M., Wong, R.O., Baylor, D.A., and Shatz, C.J. (1991). Synchronous bursts of action potentials in ganglion cells of the developing mammalian retina. Science 252, 939–943.

Meye, F.J., Soiza-Reilly, M., Smit, T., Diana, M.A., Schwarz, M.K., and Mameli, M. (2016). Shifted pallidal co-release ofGABA and glutamate in habenula drives cocaine withdrawal and relapse. Nat Neurosci 19, 1019–1024.

Miyasaka, N., Morimoto, K., Tsubokawa, T., Higashijima, S., Okamoto, H., and Yoshihara, Y. (2009). From the olfactory bulb to higher brain centers: genetic visualization of secondary olfactory pathways in zebrafish. J Neurosci 29, 4756–4767.

Mohamed, G.A., Cheng, R.K., Ho, J., Krishnan, S., Mohammad, F., Claridge-Chang, A., and Jesuthasan, S. (2017). Optical inhibition of larval zebrafish behaviour with anion channelrhodopsins. BMC Biol 15, 103.

Moorman, S.J., Burress, C., Cordova, R., and Slater, J. (1999). Stimulus dependence of the development of the zebrafish (Danio rerio) vestibular system. J Neurobiol 38, 247–258.

Moreno-Juan, V., Filipchuk, A., Anton-Bolanos, N., Mezzera, C., Gezelius, H., Andres, B., Rodriguez-Malmierca, L., Susin, R., Schaad, O., Iwasato, T., et al. (2017). Prenatal thalamic waves regulate cortical area size prior to sensory processing. Nat Commun 8, 14172.

O’Donovan, M.J. (1989). Motor activity in the isolated spinal cord of the chick embryo: synaptic drive and firing pattern of single motoneurons. J Neurosci 9, 943–958.

O’Donovan, M.J., Chub, N., and Wenner, P. (1998). Mechanisms of spontaneous activity in developing spinal networks. J Neurobiol 37, 131–145.

Okamoto, H., Agetsuma, M., and Aizawa, H. (2012). Genetic dissection of the zebrafish habenula, a possible switching board for selection of behavioral strategy to cope with fear and anxiety. Dev Neurobiol 72, 386–394.

Olstad, E.W., Ringers, C., Hansen, J.N., Wens, A., Brandt, C., Wachten, D., Yaksi, E., and Jurisch-Yaksi, N. (2019). Ciliary Beating Compartmentalizes Cerebrospinal Fluid Flow in the Brain and Regulates Ventricular Development. Curr Biol 29, 229–241 e226.

Pandey, S., Shekhar, K., Regev, A., and Schier, A.F. (2018). Comprehensive Identification and Spatial Mapping of Habenular Neuronal Types Using Single-Cell RNA-Seq. Curr Biol 28, 1052–1065 e1057.

Park, H.C., Kim, C.H., Bae, Y.K., Yeo, S.Y., Kim, S.H., Hong, S.K., Shin, J., Yoo, K.W., Hibi, M., Hirano, T., et al. (2000). Analysis of upstream elements in the HuC promoter leads to the establishment of transgenic zebrafish with fluorescent neurons. Dev Biol 227, 279–293.

Penn, A.A., and Shatz, C.J. (1999). Brain waves and brain wiring: the role of endogenous and sensory-driven neural activity in development. Pediatr Res 45, 447–458.

Reiten, I., Uslu, F.E., Fore, S., Pelgrims, R., Ringers, C., Diaz Verdugo, C., Hoffman, M., Lal, P., Kawakami, K., Pekkan, K., et al. (2017). Motile-Cilia-Mediated Flow Improves Sensitivity and Temporal Resolution of Olfactory Computations. Curr Biol 27, 166–174.

Ren, J., Qin, C., Hu, F., Tan, J., Qiu, L., Zhao, S., Feng, G., and Luo, M. (2011). Habenula “cholinergic” neurons co-release glutamate and acetylcholine and activate postsynaptic neurons via distinct transmission modes. Neuron 69, 445–452.

Romano, S.A., Perez-Schuster, V., Jouary, A., Boulanger-Weill, J., Candeo, A., Pietri, T., and Sumbre, G. (2017). An integrated calcium imaging processing toolbox for the analysis of neuronal population dynamics. PLoS Comput Biol 13, e1005526.

Rudy, J.W., Stadler-Morris, S., and Albert, P. (1987). Ontogeny of spatial navigation behaviors in the rat: dissociation of “proximal”- and “distal”-cue-based behaviors. Behav Neurosci 101, 62–73.

Satou, C., Kimura, Y., Hirata, H., Suster, M.L., Kawakami, K., and Higashijima, S. (2013). Transgenic tools to characterize neuronal properties of discrete populations of zebrafish neurons. Development 140, 3927–3931.

Scott, E.K., and Baier, H. (2009). The cellular architecture of the larval zebrafish tectum, as revealed by gal4 enhancer trap lines. Front Neural Circuits 3, 13.

Seigneur, E., Polepalli, J.S., and Sudhof, T.C. (2018). Cbln2 and Cbln4 are expressed in distinct medial habenula-interpeduncular projections and contribute to different behavioral outputs. Proc Natl Acad Sci U S A 115, E10235–E10244.

Stamatakis, A.M., Jennings, J.H., Ung, R.L., Blair, G.A., Weinberg, R.J., Neve, R.L., Boyce, F., Mattis, J., Ramakrishnan, C., Deisseroth, K., et al. (2013). A unique population of ventral tegmental area neurons inhibits the lateral habenula to promote reward. Neuron 80, 1039–1053.

Sur, M., and Leamey, C.A. (2001). Development and plasticity of cortical areas and networks. Nat Rev Neurosci 2, 251–262.

Temizer, I., Donovan, J.C., Baier, H., and Semmelhack, J.L. (2015). A Visual Pathway for Looming-Evoked Escape in Larval Zebrafish. Curr Biol 25, 1823–1834.

Tottenham, N., and Gabard-Durnam, L.J. (2017). The developing amygdala: a student of the world and a teacher of the cortex. Curr Opin Psychol 17, 55–60.

Turner, K.J., Hawkins, T.A., Yanez, J., Anadon, R., Wilson, S.W., and Folgueira, M. (2016). Afferent Connectivity of the Zebrafish Habenulae. Front Neural Circuits 10, 30.

Valente, A., Huang, K.H., Portugues, R., and Engert, F. (2012). Ontogeny of classical and operant learning behaviors in zebrafish. Learn Mem 19, 170–177.

Verdugo, C.D., Myren-Svelstad, S., Deneubourg, C., Pelgrims, R., Muto, A., Kawakami, K., Jurisch-Yaksi, N., and Yaksi, E. (2019). Glia-neuron interactions underlie state transitions to generalized seizures. bioRxiv, 509521.

Viswanath, H., Carter, A.Q., Baldwin, P.R., Molfese, D.L., and Salas, R. (2013). The medial habenula: still neglected. Front Hum Neurosci 7, 931.

Vladimirov, N., Mu, Y., Kawashima, T., Bennett, D.V., Yang, C.T., Looger, L.L., Keller, P.J., Freeman, J., and Ahrens, M.B. (2014). Light-sheet functional imaging in fictively behaving zebrafish. Nat Methods 11, 883–884.

Warden, M.R., Selimbeyoglu, A., Mirzabekov, J.J., Lo, M., Thompson, K.R., Kim, S.Y., Adhikari, A., Tye, K.M., Frank, L.M., and Deisseroth, K. (2012). A prefrontal cortex-brainstem neuronal projection that controls response to behavioural challenge. Nature 492, 428–432.

Webster, D.B., and Webster, M. (1977). Neonatal sound deprivation affects brain stem auditory nuclei. Arch Otolaryngol 103, 392–396.

White, L.E., Coppola, D.M., and Fitzpatrick, D. (2001). The contribution of sensory experience to the maturation of orientation selectivity in ferret visual cortex. Nature 411, 1049–1052.

Wiesel, T.N., and Hubel, D.H. (1965). Comparison of the effects of unilateral and bilateral eye closure on cortical unit responses in kittens. J Neurophysiol 28, 1029–1040.

Xu, H.P., Furman, M., Mineur, Y.S., Chen, H., King, S.L., Zenisek, D., Zhou, Z.J., Butts, D.A., Tian, N., Picciotto, M.R., et al. (2011). An instructive role for patterned spontaneous retinal activity in mouse visual map development. Neuron 70, 1115–1127.

Yaksi, E., Judkewitz, B., and Friedrich, R.W. (2007). Topological reorganization of odor representations in the olfactory bulb. PLoS Biol 5, e178.

Yang, Y., Cui, Y., Sang, K., Dong, Y., Ni, Z., Ma, S., and Hu, H. (2018). Ketamine blocks bursting in the lateral habenula to rapidly relieve depression. Nature 554, 317–322.

Yashina, K., Tejero-Cantero, Á., Herz, A., and Baier, H. (2019). Zebrafish exploit visual cues and geometric relationships to form a spatial memory. iScience.

Yeo, S.Y., Kim, M., Kim, H.S., Huh, T.L., and Chitnis, A.B. (2007). Fluorescent protein expression driven by her4 regulatory elements reveals the spatiotemporal pattern of Notch signaling in the nervous system of zebrafish embryos. Dev Biol 301, 555–567.

Yu, Y.C., He, S., Chen, S., Fu, Y., Brown, K.N., Yao, X.H., Ma, J., Gao, K.P., Sosinsky, G.E., Huang, K., et al. (2012). Preferential electrical coupling regulates neocortical lineage-dependent microcircuit assembly. Nature 486, 113–117.

Yuste, R. (1997). Introduction: spontaneous activity in the developing central nervous system. Semin Cell Dev Biol 8, 1–4.

Yuste, R., Peinado, A., and Katz, L.C. (1992). Neuronal domains in developing neocortex. Science 257, 665–669.

Zhang, B.B., Yao, Y.Y., Zhang, H.F., Kawakami, K., and Du, J.L. (2017). Left Habenula Mediates Light-Preference Behavior in Zebrafish via an Asymmetrical Visual Pathway. Neuron 93, 914–928 e914.

Zhang, R.W., Du, W.J., Prober, D.A., and Du, J.L. (2019). Muller Glial Cells Participate in Retinal Waves via Glutamate Transporters and AMPA Receptors. Cell Rep 27, 2871–2880 e2872.

Zou, D.J., Feinstein, P., Rivers, A.L., Mathews, G.A., Kim, A., Greer, C.A., Mombaerts, P., and Firestein, S. (2004). Postnatal refinement of peripheral olfactory projections. Science 304, 1976–1979.

Wetherington, J., Serrano, G., and Dingledine, R. (2008). Astrocytes in the epileptic brain. Neuron 58, 168–178.

Wyatt, C., Bartoszek, E.M., and Yaksi, E. (2015). Methods for studying the zebrafish brain: past, present and future. Eur J Neurosci 42, 1746–1763.

Yasuda, C.L., Chen, Z., Beltramini, G.C., Coan, A.C., Morita, M.E., Kubota, B., Bergo, F., Beaulieu, C., Cendes, F., and Gross, D.W. (2015). Aberrant topological patterns of brain structural network in temporal lobe epilepsy. Epilepsia 56, 1992–2002.

